# ColiCoords: A Python package for the analysis of bacterial fluorescence microscopy data

**DOI:** 10.1101/608109

**Authors:** Jochem H. Smit, Yichen Li, Eliza M. Warszawik, Andreas Herrmann, Thorben Cordes

## Abstract

Single-molecule fluorescence microscopy studies of bacteria provide unique insights into the mechanisms of cellular processes and protein machineries in ways that are unrivalled by any other technique. With the cost of microscopes dropping and the availability of fully automated microscopes, the volume of microscopy data produced has increased tremendously. These developments have moved the bottleneck of throughput from image acquisition and sample preparation to data analysis. Furthermore, requirements for analysis procedures have become more stringent given the requirement of various journals to make data and analysis procedures available. To address this we have developed a new data analysis package for analysis of fluorescence microscopy data of rod-like cells. Our software ColiCoords structures microscopy data at the single-cell level and implements a coordinate system describing each cell. This allows for the transformation of Cartesian coordinates of both cellular images (e.g. from transmission light or fluorescence microscopy) and single-molecule localization microscopy (SMLM) data to cellular coordinates. Using this transformation, many cells can be combined to increase the statistical significance of fluorescence microscopy datasets of any kind. Coli-Coords is open source, implemented in the programming language Python, and is extensively documented. This allows for modifications for specific needs or to inspect and publish data analysis procedures. By providing a format that allows for easy sharing of code and associated data, we intend to promote open and reproducible research.

The source code and documentation can be found via the project’s GitHub page.

## Introduction

Fluorescence microscopy has become a crucial tool in studying bacterial cell biology(1–8). It is minimally invasive and allows for the study of living bacteria in a controlled environment and to monitor the motion and sub-cellular topologies of any proteinaceous factor(9–24) or nucleic acids(25–28). Through the multitude of genetically programmable fluorescent protein probes(29–36) and commercially available dyes(37–41) with different conjugation capabilities(42–46), virtually all components of the bacterial machinery can be studied with high specificity and spatio-temporal resolution. Even the detection of single fluorescent probes has become a routine experiment, revealing both dynamics and heterogeneity.

However, with the introduction of fully automated(47) and autonomously operating microscopes(48, 49), images can be acquired independently of experimentalist interference at high acquisition rates(50–52). A typical experiment conducted overnight spanning the dimensions of position, time and channels can easily generate thousands of images. This explosion of available multidimensional data has made analysis the new bottleneck in single-cell fluorescence microscopy studies. Furthermore, in the spirit of reproducibility and open availability of scientific results, more and more peer-reviewed journals require deposition of source data and evaluation methods – a process that is hampered by non-standardized evaluation routines and lab-specific software, often based on commercial platforms.

The exception is current options for bacterial image analysis such as SuperSegger(62, 63), Oufti(64) and MicrobeJ(65). Although these data analysis tools are able to tackle a lot of the problems inherent to (bacterial) live-cell image analysis, we have identified several drawbacks which we aimed to address with ColiCoords. The main points are related to the platform, structure and philosophy of the data analysis routine rather than the functionality for analysing images.

First, with available tools, it is difficult for users themselves to either inspect the exact mathematical procedures applied to their data or customize them. The source code of these analysis packages are freely available and licensed under the GNU General Public License. Modification of Oufti and SuperSegger, however, require a Matlab license, which is not available in many institutions. MicrobeJ, on the other hand, can be freely modified, but users face a daunting task due to the shear size of the project and the lack of docstrings.

ColiCoords is written in the freely available language Python and the source code has been released under the MIT license. The code is available on GitHub and is extensively documented with both docstrings, online documentation and its basic principles are described in this paper. Being hosted on GitHub together with Continuous Integration for testing makes ColiCoords an ideal platform for other users to contribute and modify its code.

Second, most other currently available analysis options based on a graphical user interface (GUI). We would argue here that GUIs are inherently limiting and obfuscate the exact data analysis procedure. While GUI-based analysis is intuitive it limits the use to the available graphical elements, with little flexibility. Importantly, due to the large number of permutations to the of order of operations that can be executed via a GUI, the exact data analysis procedure is typically not documented in detail and thus hard to reproduce.

In contrast, ColiCoords offers a basic foundation for single cell data analysis where users can apply all functionality from Python itself, the SciPy(66) ecosystem and community developed image analysis procedures. The workflow of Coli-Coords is based on Jupyter Notebooks(67, 68), which act as a lab journal page, where all steps of data analysis and its associated parameters can be executed and described and results are plotted in interactive graphs. This type of interactive analysis workflow is being adopted rapidly in the scientific community(69). An example from the microscopy community is the smFRET analysis package FRETBursts(70).

In addition, ColiCoords is open-ended, meaning it can be used in a pipeline of analysis where users can freely choose pre- and post-processing options (Figure 1). The input data can be either image-based (brightfield, phase contract, fluorescence) or sparse data (super-resolution localizations). Users are therefore free to choose preprocessing options for tasks such as segmentation(53, 54, 56, 57, 64) and super-resolution reconstruction(58–61). Colicoords is out of the box compatible with High-performance computing (HPC) as operations on a per-cell basis can be computed in parallel.

**Fig. 1.**
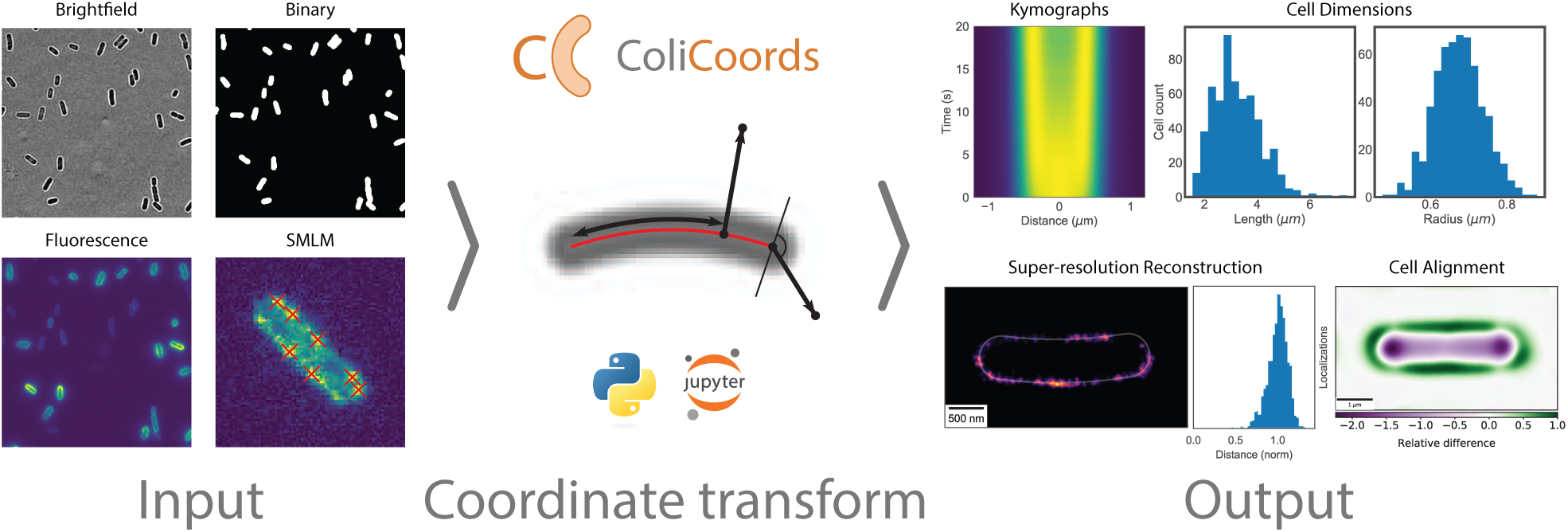
Overview of workflow pipeline using ColiCoords. Input data is either image data or sparse data (localizations). Input images need to be segmented to identify cell location and orientation. Third party options for image segmentation include Ilastik(53), CellProfiler(54, 55) or Keras(56)/Tensorflow(57). SMLM data have to be reconstructed by external software such as DAOSTORM(58, 59), ThunderSTORM(60), QuickPALM(61) or others prior to use. ColiCoords can then be used to transform the Cartesian coordinates of the input data to cellular coordinates. The transformed data can to generate output graphs, such as kymographs, histograms of the cell’s dimensions, axial distributions or to align cells.

The central feature of bacterial image analysis in ColiCoords is transforming input Cartesian coordinates (image pixel co-ordinates or super-resolution localizations) to cellular coordinates. The coordinate system can be derived from different sources, including binary images, brightfield images or even super-resolution membrane markers. With the coordinate system in place, ColiCoords offers several different visualization and analysis methods, including alignment of cells, radial, longitudinal or angular (poles) distributions, kymographs and super-resolution reconstruction. (Figure 1).

## ColiCoords principles and features

### A. Coordinate definitions

The general principle of the coordinate system is shown in Figure 2. Here, a brightfield image of an *E. coli* cell is shown with the coordinate system overlayed. By establishing a per-cell coordinate system every pair of Cartesian coordinates (*x*_*p*_, *y*_*p*_) from either pixels or fluorescent foci can be mapped to cell coordinates (*l*_*c*_, *r*_*c*_, *ϕ*); Figure 2**A**. The transformation is based on a second-degree polynomial (*p*(*x*), *x*_*l*_ ≤ *x* ≤ *x*_*r*_) (red line) which corresponds to the midline of the cell. The parameters describing *p*(*x*) are initially based on guess values which can be refined after. To transform a point (*x*_*p*_, *y*_*p*_), the first step is to find the point (*x*_*c*_, *y*_*c*_) on *p*(*x*) which is closest to (*x*_*p*_, *y*_*p*_), perpendicular to *p*(*x*). This is done by minimizing the squared distance between both points:

**Fig. 2.**
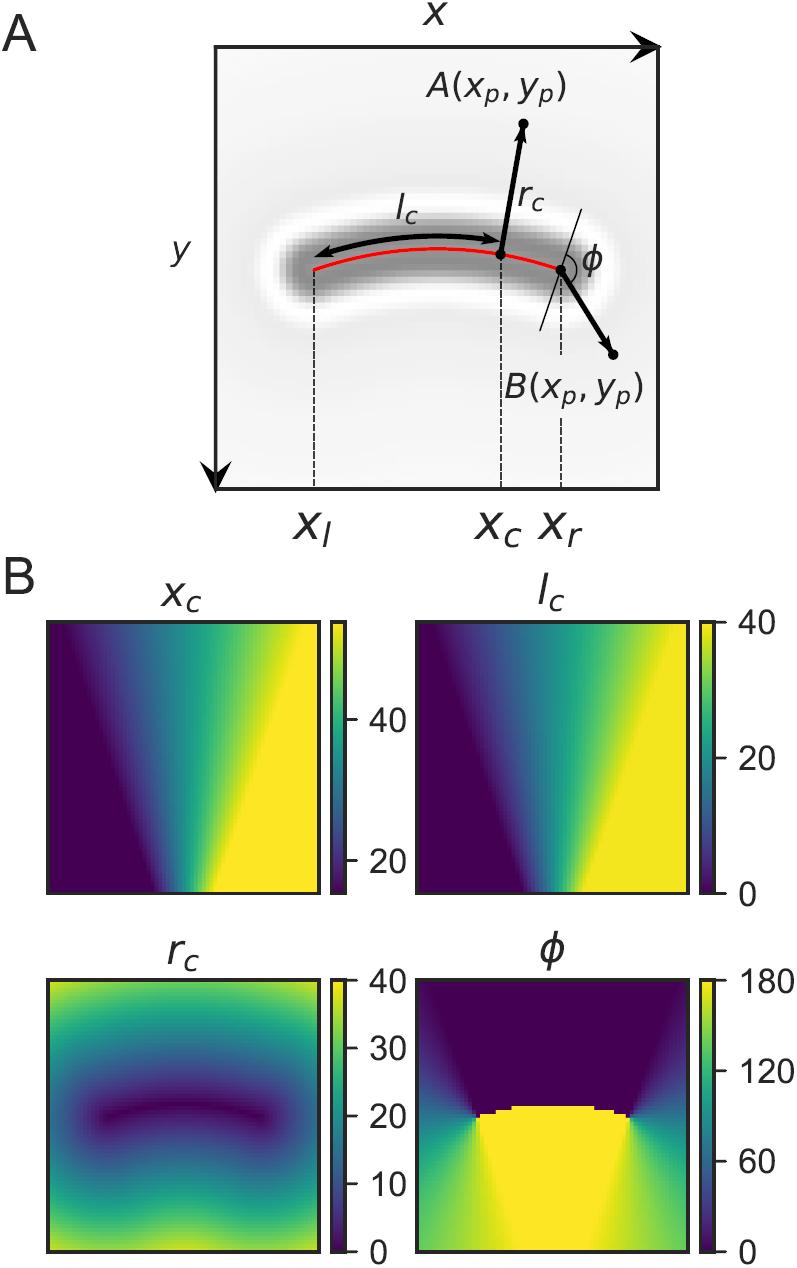
General description of the in-cell coordinate system. **A** Brightfield image of an *E. coli* cell with coordinate system overlayed. Every point with coordinates *x*_*p*_, *y*_*p*_ can be transformed to coordinates *l*_*c*_, *r*_*c*_, *ϕ*. **B** Images showing the values of *x*_*c*_ as well as cellular coordinates values *l*_*c*_, *r*_*c*_, *ϕ*.

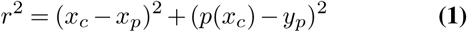

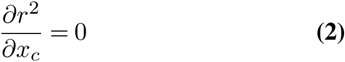

Solving equation 2 gives a cubic equation which is solved analytically to ensure fast transformation of many data points. To account for points B at the poles of the cells the coordinate *x*_*c*_ is restricted to the domain [*x*_*l*_, *x*_*r*_]. The cellular coordinates (*l*_*c*_, *r*_*c*_, *ϕ*) can then be calculated from *x*_*c*_ and *p*(*x*). The longitudinal coordinate *l*_*c*_ is given by the arc length along *p*(*x*) from *x*_*l*_ to *x*_*c*_:

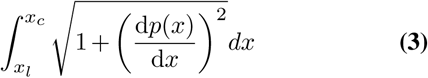

This expression also gives the full length of the cell when *x*_*c*_ is substituted by *x*_*r*_. The radial coordinate *r*_*c*_ is simply the euclidean distance between (*x*_*c*_, *p*(*x*_*c*_)) and (*x*_*p*_, *y*_*p*_):

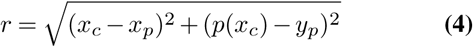

The third coordinate is the angle *ϕ* (in degrees) and it uniquely defines points A along the body of the cell by distinguishing between the top and bottom of the cell, as well as defining the position of points B at the poles. The value of *ϕ* is 0 at the top of the cell runs from 0 to 180 along the right pole, where *ϕ* is defined as the angle between the line perpendicular to *p*(*x*) at *x*_*r*_ and the line from (*x*_*c*_, *p*(*x*_*c*_)) to (*x*_*p*_, *y*_*p*_). For points below the midline of the cell the value of *ϕ* is 180, which then runs from 180 back to 0 along the left pole.

Note that the top area of the cell, as it is displayed in Panel **A** of Figure 2, is given by *y*_*p*_ *< p*(*x*_*p*_), since the origin of the Cartesian coordinates is at the top left, where the postive y-axis is running down. By this definition, the coordinate of the midpoint of the top-left pixel is (0.5, 0.5), in line with the coordinate definitions in the image analysis software ImageJ(71, 72).

### B. Preprocessing and optimization

To analyse fluorescence microscopy data with ColiCoords some preprocessing of the input data is required. Notably, ColiCoords is not designed to segment or detect cells in the raw microscopy images. Binary (labelled) input images are required for Coli-Coords processing to identify the location of cells in the images. However, these binaries are only needed to localize cells and initialize the coordinate system, and they are not needed to form the final optimized coordinate system, consequently a high accuracy of segmentation is not required. Binary or segmented images can be derived from brightfield, differential interference contrast (DIC), phase contrast or fluorescence images by established tools such as CellProfiler (54), Ilastik(53), Oufti(64) or Convolutional Neural Networks (e.g. implemented by TensorFlow(57) or Keras(56)). An implementation of the U-Net segmentation architecture(73) is included in ColiCoords together with examples on how to use it.

Fluorescence images should be (optionally) background or illumination corrected and different channels such as brightfield or DIC should be aligned if they are acquired with different optics. ColiCoords allows the processing of any kind of image data as well as sparse data, such as single-molecule tracking and single-molecule localization microscopy (SMLM) data. Based on the provided binary images, single-cells are automatically cut out from all image data, the positions of the localizations are selected and transferred to the coordinates of the cropped image. The orientation of the cell is calculated using the binary image by calculating the image moments(74). All data elements are then rotated to orient the cells horizontally. The result of this process is a collection of single-cell Python-objects where all data are organized on a cell-by-cell basis, and every cell has its own coordinate system which can be used to perform calculations, analysis or visualizations on the data element of choice. These sets of cells can then be indexed and selected based on user-defined criteria (shape, size, fluorescent signal etc.) and further analysed or inspected through interactive plots within the Jupyter Notebook environment.

After initialization of the cell objects, the coordinate system needs to be refined to allow it to more accurately describe the cell’s shape. ColiCoords implements the optimization of the coordinate system based on different data sources by providing several objective functions which, when minimized, give the best matching coordinate system for the respective data source. Any image-type data (e.g. brightfield) can be used provided there is an anisotropic signal along the length of the cell, as well as localization (SMLM) data provided the localizations describe the outline of the cell.

Figure 3 shows the iterative optimization process. In Panel **A** the ground truth binary image (derived from measurement) is shown together with the cell’s coordinate system. The coordinate system is described by a total of 6 parameters, 3 coefficients *a*_0_, *a*_1_, *a*_2_ which define the second degree polynomial *p*(*x*), the left and right endpoints *x*_*l*_ and *x*_*r*_, and the cell’s radius *r*. In the figure, the polynomial *p*(*x*) is shown together with the isodistance line from *p*(*x*) with distance *r*. As can be seen from the figure, the initial coordinate system does not accurately describe the cell’s shape. To refine the coordinate system, first a radial distance image is calculated (Panel **B**). This image is then thresholded with the initial guess value of *r*, which gives a new binary image (Panel **C**, black). Comparison of this image with the ground truth binary image gives the *χ*^2^ value, which is minimized to optimize the fit of the coordinate system. The fitting in ColiCoords is done via the package Symfit(75), which provides an API to the minimizers implemented by `scipy.minimize`(66).

**Fig. 3.**
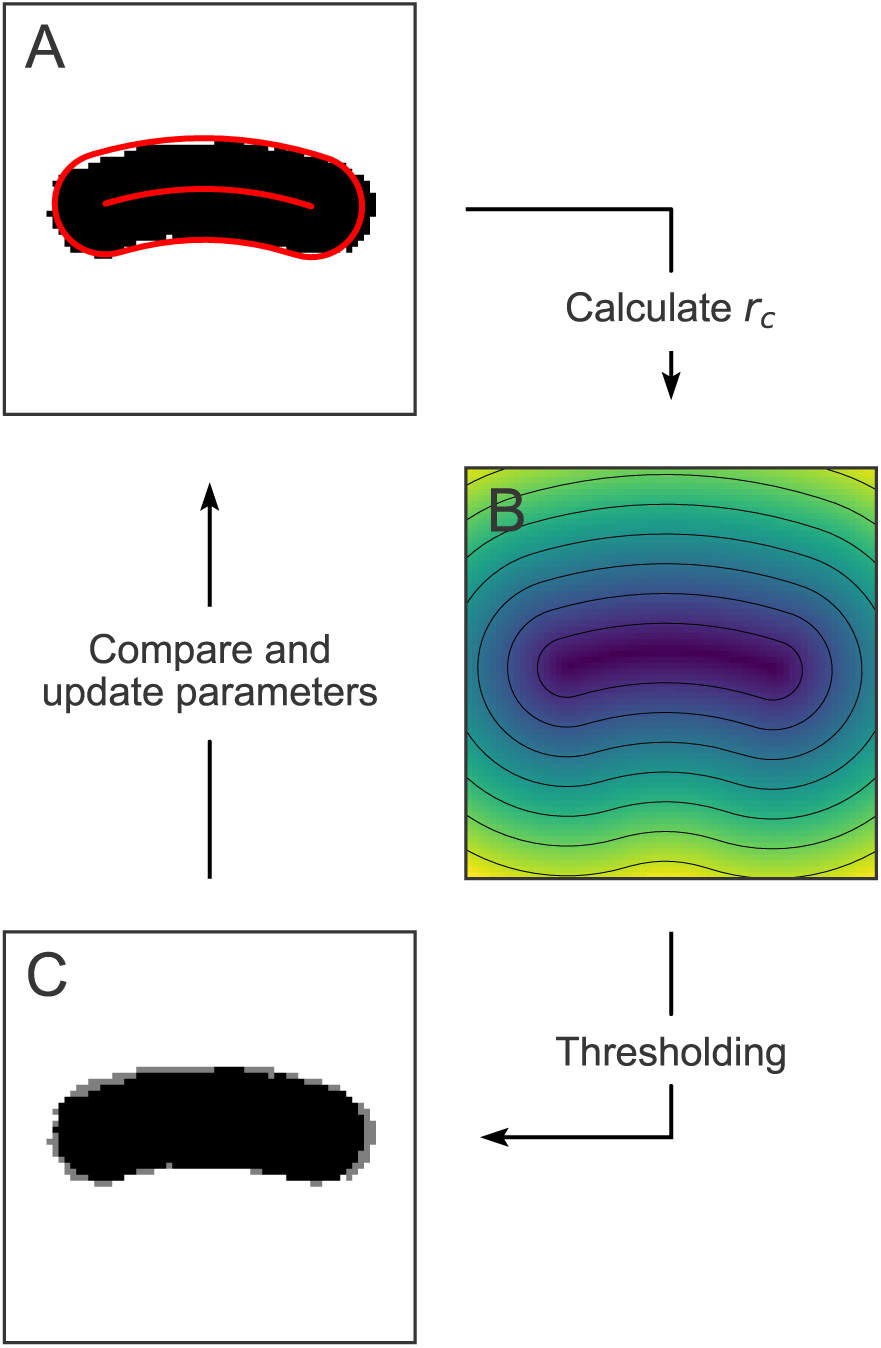
Optimization of the cell’s internal coordinate system. **A** Ground-truth binary image with cell midline and outline calculated from initial guess parameters. **B** Radial distance image calculated from initial guess coordinates. **C** Calculated binary image (black) obtained by thresholding the distance image superimposed on the ground truth image (grey).

Other microscopy images can be used to optimize the coordinate system, where the only constraint is that the signal needs to be isotropic along the length of the cell. This is shown for binary, brightfield and fluorescence images in Figure 4. The figure shows the three different channels measured for a single cell. The binary is derived from the brightfield and is used for initial guesses of the coordinate system. This initial guess coordinate system is shown in red in the left column. This coordinate system is used to calculate a synthetic image, averaged along the angular coordinate (middle). A comparison between this image and the measurement gives a measure for how well the current parameters describe the shape of the cell. The optimal parameters can be found via iterative optimization (right). For brightfield- or fluorescence data-based optimization the radius *r* of the cell is determined in a second step by determining the half-maximum point of pixel intensity.

**Fig. 4.**
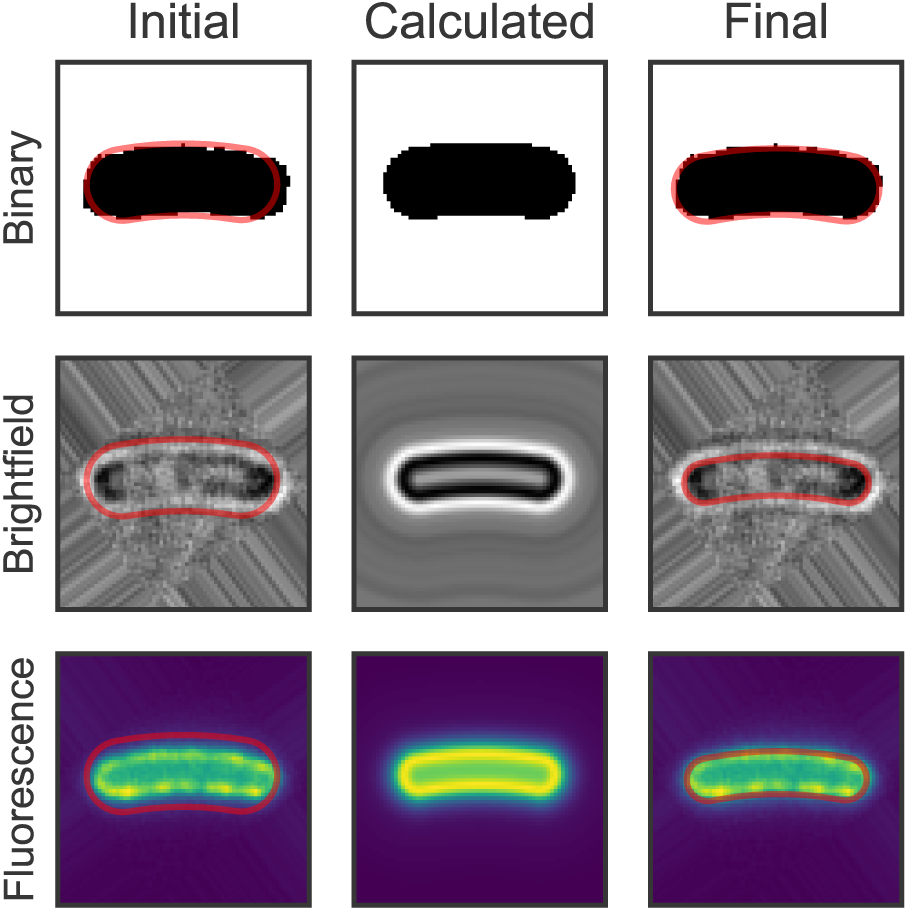
Iterative optimization of the coordinate system based on binary, brightfield and fluorescence images.

Finally, ColiCoords can optimize the coordinate system based on SMLM data of a membrane marker. This type of data can be obtained through various super-resolution techniques such as Point Accumulation for Imaging in Nanoscale Topography (PAINT)(27, 76–78), Stochastic Optical Reconstruction Microscopy (STORM)(79, 80) or PhotoActivated Localization Microscopy (PALM)(81, 82) imaging.

The optimization process and a possible application thereof is illustrated in Figure 5. Panel **A** shows a STORM super-resolution reconstruction of the membrane protein marker LacY-eYFP. In the top panels, the coordinate system based on initial guesses (derived from binary image; grey) is shown. The bottom panel shows the result upon optimization based on the STORM-localizations data. Here, the radial distance *r*_*c*_ for all localizations is calculated and by comparison to the value for *r* the *χ*^2^ (squared differences) is calculated. Minimization of the *χ*^2^ gives the parameters where the coordinate system that most appropriately describes the bacterial membrane. Interestingly, the localizations along the membrane show a periodic fine structure. To further characterize this, the position along the perimeter of the cell was plotted in panel **B**. The zero-point position is defined as the beginning of the ‘top’ part of the membrane (*l*_*c*_ = 0, *ϕ* = 0, *r*_*c*_ = *r*, panel **A**) and the positive directions runs clockwise along the cell outline.

**Fig. 5.**
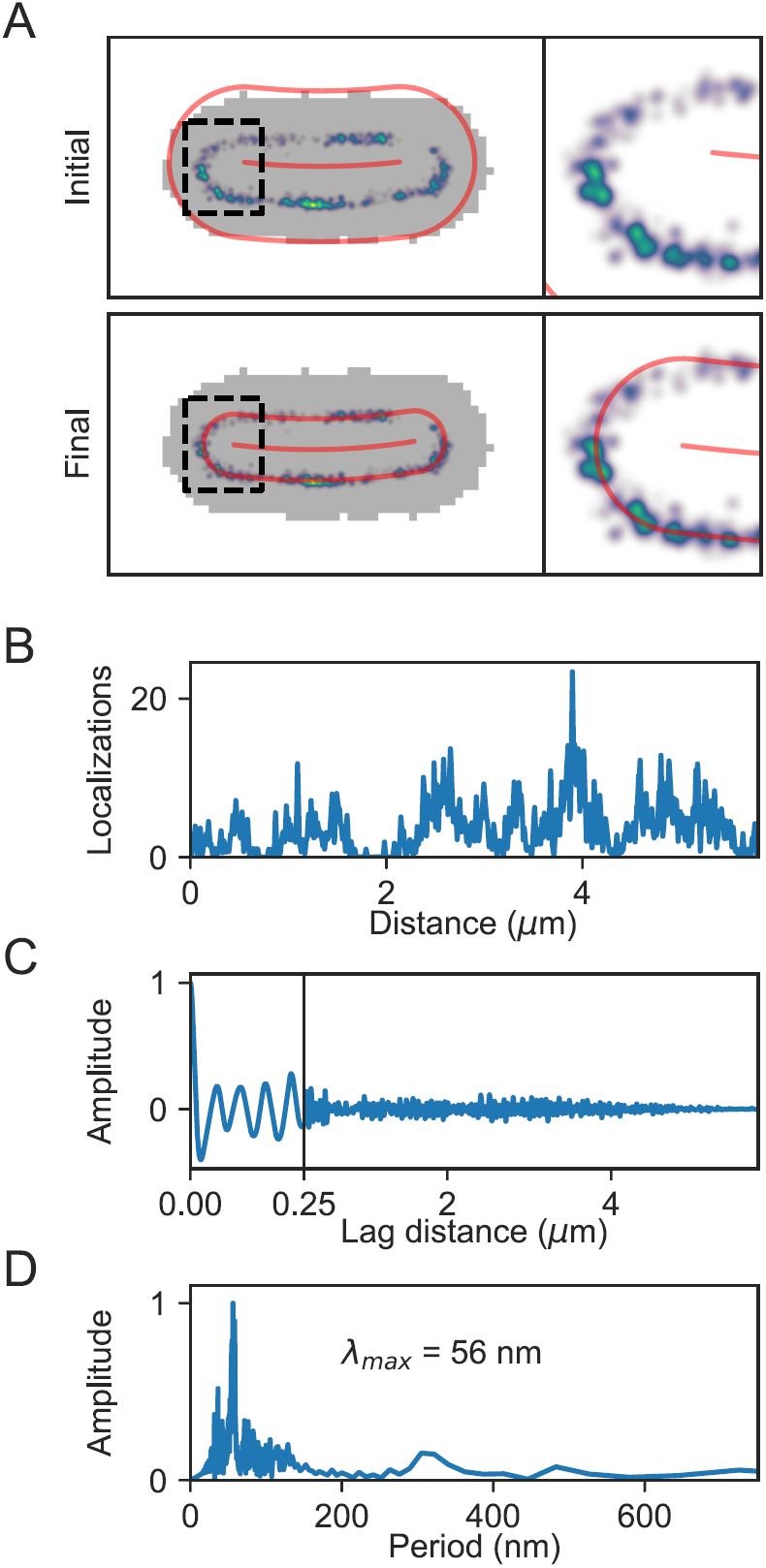
Iterative optimization of the coordinate system based STORM superresolution data of the membrane marker LacY-eYFP. **A** Top: STORM reconstruction on top of the *E. coli* binary image (grey) with initial guesses coordinate system in red. Bottom: Coordinate system after optimization based on STORM localizations. **B** STORM reconstruction along the perimeter of the cell showing localizations as a function of distance along the membrane. **C** Spatial autocorrelation function of **B**. **D** Fourier transform of **C** showing the largest amplitude at a periodicity of 56 nm.

To extract the size of the periodic structures observed, at first a spatial auto-correlation(83) function was calculated and its low-frequency components were subtracted by means of a sliding window, to reveal clear oscillations (panel **C**). By Fourier transforming this signal (Panel **D**) we found a periodicity of the signal of 52 nm, which we attribute to an artefact resulting from the YFP fusion, as previously reported for MreB-YFP constructs(84).

In conclusion, optimization based on a SMLM super-resolution membrane marker is expected to provide the most accurate approach for creating a coordinate system within the cell (vide infra, section **D**). This allows for aligning and combination of multiplexed localization data from many cells with high accuracy.

### C. Batch processing

To demonstrate the ability of Coli-Coords to analyse a datasets consisting of many cells, we applied a typical analysis procedure to two sets of fluorescence microscopy images. The first set of *E. coli* cells were incubated with the reactive dye Cy3B-NHS to label the outer membrane, while the second set of cells expressed eGFP in the cytosol.

For each condition, a subset of 100 images were manually annotated through the use of a custom GUI element. These annotated binary images were then combined and used to train a convolutional neural network. An implementation of the U-Net architecture(73) neural network was used, based on the Keras(56) API using Tensorflow(57) as backend.

The brightfield images were first scaled down to 256×256 to preserve graphics card memory. A custom implementation of Keras’ `Sequence` was used for preprocessing and augmentation. All input images were normalized using tanh-estimators(85) and augmented 8-fold through permutations of horizontal mirroring, vertical mirroring and transposing the images. This augmented data was randomly split into a set validation data and training data.

After training the neural network, the network was applied to the whole set of images to generate binary images. The images were then scaled back to 512×512 pixels. Next, binary shapes were filtered from the segmentation mask by their shape and size. The resulting binary images were used as input for ColiCoords, together with the corresponding bright-field and fluorescence images. The cell objects were filtered based on the results of optimizations by binary images and brightfield images as well as a measurement of the cell’s radius from the brightfield image. This yielded a total of 2341 and 1691 cells for the Cy3B and eGFP datasets, respectively. The analysis output is shown in Figure 6. In panel **A**, two individual cells for each condition are shown. Their radial distribution profile is shown in panel **B** (blue line), together with the individual datapoints from each pixel as red points. In panel **C**, all cells from each dataset were aligned by transforming all pixel coordinates in the fluorescence images from each cell to cellular coordinates. These cellular coordinates are then transformed back onto a model cell with user-specified dimensions. The resulting point cloud is then used to generate an aligned fluorescence image by convolution with a 2D gaussian. The final is image is displayed with spline interpolation. In panel **D**, the average radial distribution for all cells is shown. The curves are individually normalized in the x-direction by setting the brightfield-measured radius to 1 whereas the y-direction is normalized by the maximum of each curve. The shaded region (standard deviation) therefore only reports on variations in the shape of the radial distribution profile.

**Fig. 6.**
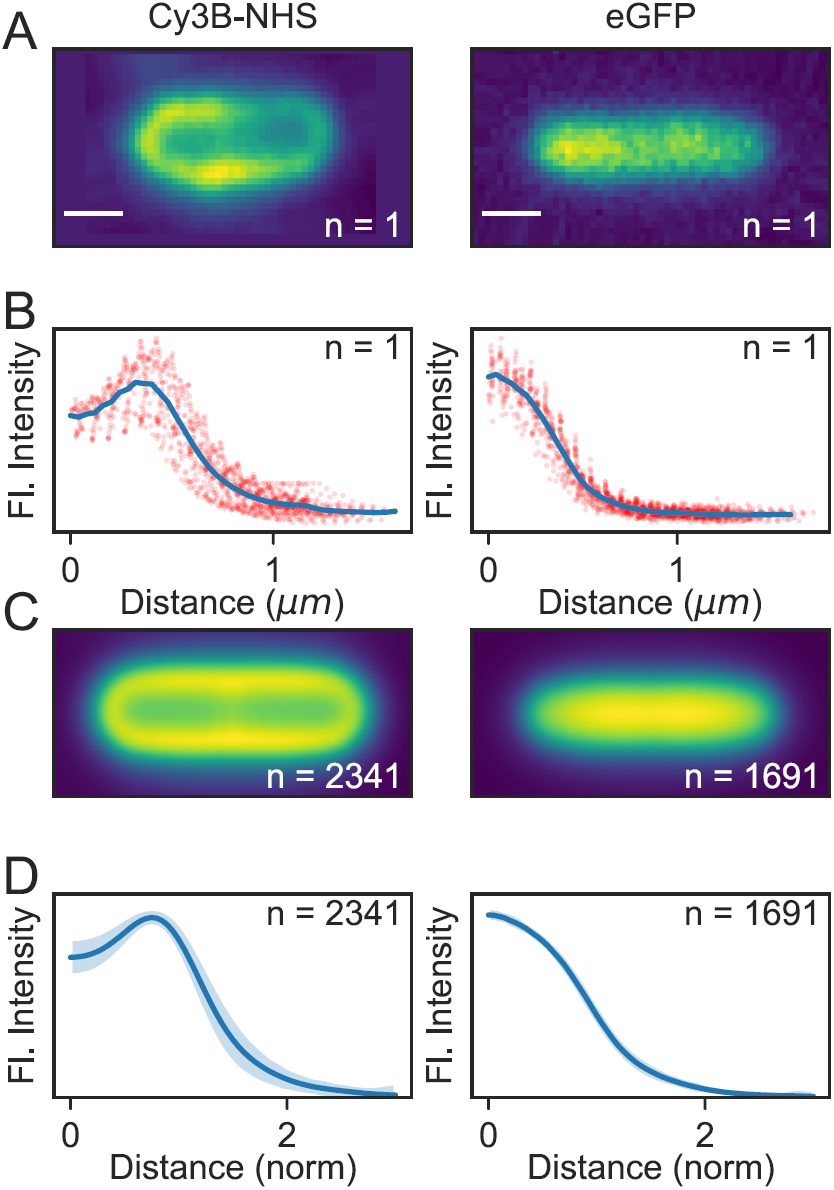
Batch processing of several thousand cell objects. Two datasets were analyzed, one where cells are labelled on the outer membrane with Cy3B-NHS and one with cell expressing eGFP in the cytosol. **A** Fluorescence images of single *E. coli* cells, scale bar 750 nm. **B** Radial distribution profiles for the cells in **A**. Individual datapoints from every pixel in the image are shown as red points, together with the resulting radial distribution (blue line). **C** Aligned and average fluorescence images for n=2341 and n=1691 cells, respectively. **D** Average radial distribution profiles. The standard deviation is shown as a shaded region.

### D. Synthetic benchmarks

In order to evaluate Coli-Coords’ performance a synthetic dataset was generated and subsequently analyzed. By comparing the results for different conditions and optimization methods the accuracy can be benchmarked.

First, a set of 10000 synthetic cells were generated using ColiCoords’ `synthetic_data` module. The geometric parameters describing these synthetic cells where chosen based on a set of measured *E. coli* cells. These synthetic cells have a binary image data element by default. A brightfield data element was added based on normalized brightfield radial distributions measured from *E. coli*. The ratio between the background intensity (no cells) of the brightfield and the maximum of the scattered light from the bacterial membrane was measured and kept constant to replicate realistic signal-to-noise ratios. The background light was set to one and then multiplied by the chosen photon count number (500, 1000 and 10000), after which the resulting values where drawn from a Poisson distribution to simulate shot noise. Finally, normally distributed noise was added with a standard deviation of 20 photon counts.

To each cell two homogeneously distributed membrane-localized STORM data elements were added. The first data element describes localizations at the inner membrane. The mean radial distance of the localizations was set according to where the inner membrane should be with respect to the radius of the cell measured from the brightfield image based on measurements with the construct LacY-eYFP. The standard deviation from the mean radius was set to 0.25 pixels (20 nm). A second STORM data element was added 100 nm further out with the same standard deviation.

From this set of synthetic cells, images where reconstructed. An average of 10 ± 3 cells where taken from the set and randomly rotated and placed in each image of 512×512 pixels (40 *µ*m × 40 *µ*m), ensuring at least 5 pixels distance between the cells’ binary images. The coordinates of the cell’s STORM data element where combined into one big STORM localizations table.

These processes yielded 1000 ground-truth binary images together with 3 sets of 1000 brightfield images with different photon numbers (500, 1000, 10000 per pixel) and therefore different signal-to-noise ratios (Example images in Figure 7**A**). The whole ColiCoords’ data analysis pipeline was performed on this dataset to benchmark its performance.

**Fig. 7.**
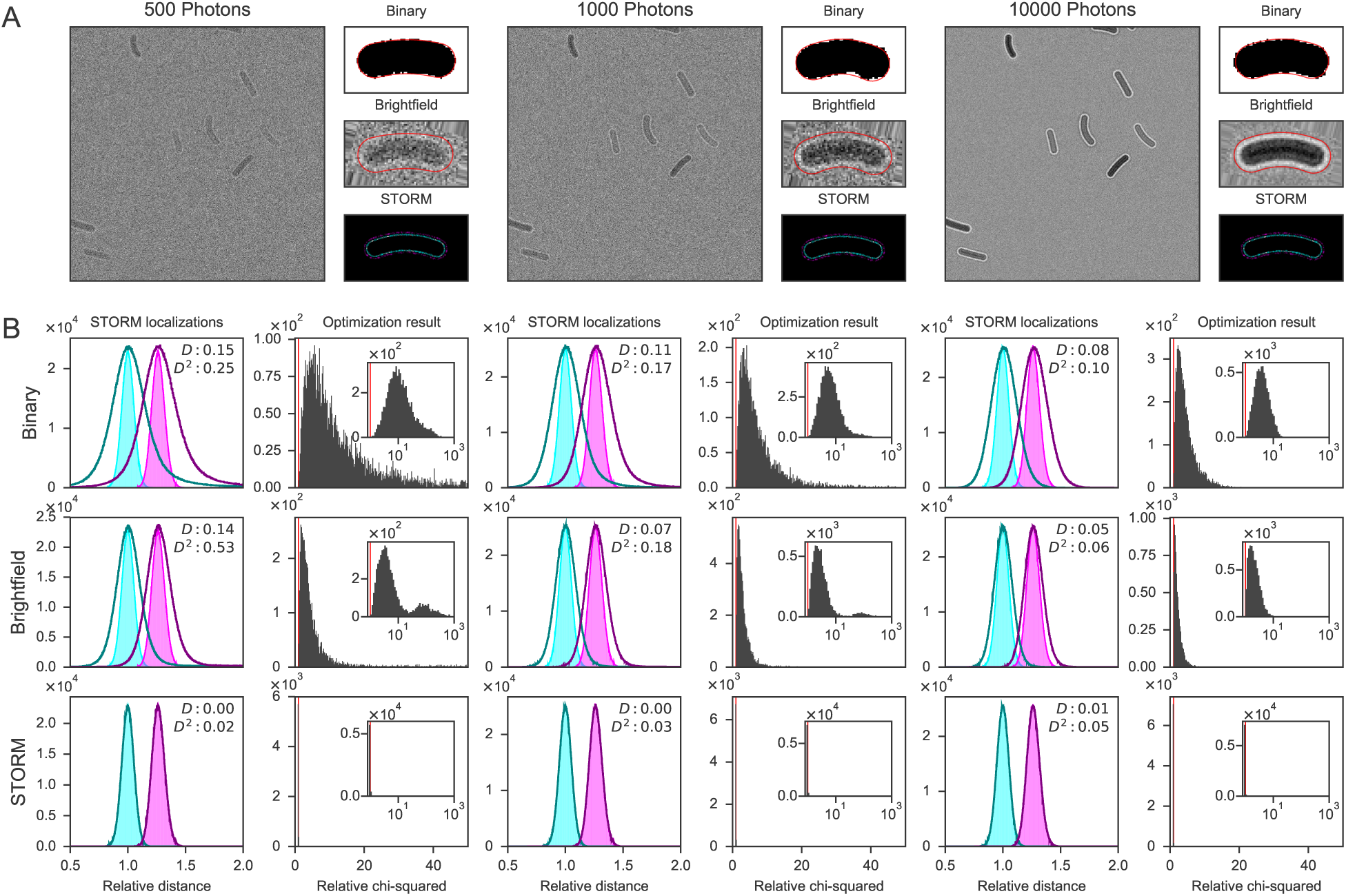
Benchmarking of ColiCoords software with a synthetic dataset. **A** Examples of generated input data brightfield images for different photon counts (500, 1000 and 10000 average per pixel) and example of one processed single cell. Different data elements (Binary, Brightfield, STORM) are shown together with the outline of the coordinate system (red/white line) where the coordinate system is optimized based on that data element. **B** Evaluation of the coordinate system of n=6569, 7127, 7245 cells out of 7245. Left: relative radial distance of all inner (cyan) and outer (magenta) membrane STORM localizations. The radial position is normalized to the ground-truth inner membrane position. The ground-truth radial distance (light colours, filled histogram) is compared with radial distances calculated with the coordinate system as calculated for the different data elements (binary, brightfield, STORM) for different brightfield image photon counts (dark colours, line only). The absolute mean deviation (*D*) and the root mean squared deviation (*D*^2^) are shown in the graph. Right: Relative minimization objective function *χ*^2^. The *χ*^2^ is calculated for the obtained coordinate system for each cell for the STORM data element for each condition. The obtained value is divided by the ground-truth *χ*^2^ value, therefore a value of 1 indices a perfectly fitted coordinate system (red line).

First, the brightfield images were segmented. For all 3 signal-to-noise conditions, the first 400 images where used to train the neural network. After training, the network was applied to the whole set of 1000 images to generate binary images.

Next, the binary images were filtered. Bordering cells were removed and the remaining binary objects were selected based on their size and ellipticity where the selection criteria were obtained from the ground-truth binary images. The filtered and labelled binary images were then used by Coli-Coords together with the brightfield and STORM data to generate a set of ‘measured’ cell objects.

The ‘measured’ cell objects obtained are correlated to the original ground-truth cell objects by first performing a cluster analysis on the input STORM table(86). The ground-truth cells are then identified by encoding the combined STORM intensity and comparing to the ground-truth intensity of the STORM clusters. The corresponding measured cell was then found by comparing the centre-of-mass positions of the STORM clusters and the filtered binary images used as input. This yielded two sets of cells, one with ground-truth cells and one with corresponding ‘measured’ cells for comparison.

Cells which had too few STORM localizations compared to their ground-truth counterparts were discarded due to them being too poorly segmented. Several cells had too many STORM localizations due to localizations from neighbouring cells. These additional localizations were also removed. This yielded a total of 6569, 7129 and 7245 cells out of maximum 7245 cells for brightfield images with 500, 1000 and 10000 photons, respectively. This number is lower for input images with lower signal-to-noise because some cells are segmented poorly by the neural network.

Finally, the coordinate system of the cells was optimized using different approaches to compare their accuracy. The cells in each condition were initialized with guesses derived from segmented binary images and subsequently optimized based on the data elements binary, brightfield and storm (inner membrane). This yielded a total of 9 sets of cells which are summarized in panel **B** of Figure 7. In the left column of each condition the relative radial distances of all STORM localizations in the dataset are plotted as calculated by its coordinate system. The radial distances are normalized to the ground truth inner membrane distance to eliminate cell-to-cell radius variations. The ground truth radial distances of the inner (cyan) and outer (magenta) are shown in light colours and filled histogram. The calculated radial distances are shown on top with dark colours where the outer line of the histogram is shown only.

From these graphs the effect of the accuracy of the coordinate system can be shown on the resulting radial distribution graph. As can be seen, in the condition with the lowest number of photons (500) and a coordinate system optimized on the binary data element, the resulting radial distribution histogram is significantly wider than the ground-truth radial distribution. Compared to binary images derived from higher signal-to-noise images, the result more accurately describes the underlying ground-truth. This is further reflected by comparing the obtained *χ*^2^ values from the optimization process. In the histogram on the right (second column) the relative *χ*^2^ of each cell derived from its optimized coordinate system is plotted. The values are obtained by calculating the *χ*^2^ value for STORM optimization (mean squared difference in distance between coordinate system outline and STORM localizations) and dividing this value by the ground-truth *χ*^2^ value to get a relative *χ*^2^. This value is an indication for the goodness of the fit, where a value of one indicates a perfect fit (red line in histograms).

As can be seen in the histogram, for 500 photons and optimization based on the binary image (Panel **B**, top left), the median *χ*^2^ value of optimization is roughly 10-fold higher compared to the ground-truth values. The inset (log scale x-axis and logarithmically increasing bin size) shows a long tail in the distribution to higher relative *χ*^2^ values up to 500.

When comparing the binary optimization result for 1000 and 10000 photons, it can be seen that the radial distribution histograms become narrower and more accurately match the ground truth radial distribution. The distribution of relative *χ*^2^ values also reflect this with median values of 5.8 an 3.8, respectively.

Moving one row down the results are plotted for optimization based on the brightfield image. The initial guesses for optimization derived from the binary images and therefore the quality of the segmentation influences the final result. As can be seen from the radial distribution histograms, the coordinate system obtained more accurately describes the cell compared to binary optimization. For 10000 photons the radial distribution graph very closely matches the ground truth. In the relative *χ*^2^ distributions a second population can be seen which is mostly present for lower photon numbers. This is a consequence of poor initial guesses and the optimization therefore returns a different local minimum. The median relative *χ*^2^ values are 4.1, 2.6 and 1.9, respectively.

The third row shows the result from optimization based on the inner membrane STORM localization dataset. Since the STORM localizations are identical in all cells and are independent of the brightfield image, any difference in the optimization result is due to the initial guesses. As expected, the obtained radial distribution histograms almost perfectly overlap with the ground truth histograms. The obtained median relative *χ*^2^ values are all 0.99, however some small outliers are found, where the population sizes of values >50 are 2, 3 and 9 respectively.

In Figure 7 for 10000 photons and brightfield optimization it can be seen that the ground-truth result is matched closely but not completely. This is unexpected since the cells were generated from a perfect second-degree polynomial shape and therefore it was expected that the optimization result would more accurately describe the cell’s shape. The *χ*^2^ value of the obtained result was lower compared to the ground-truth, thus the problem is not that the algorithm returned a local minimum instead of a global minimum. Therefore, switching to a global optimization algorithm such as differential evolution(87) will not yield the desired improved result.

Instead, it was found that in the optimizer’s implementation a choice of speed over accuracy is the origin of the discrepancy. Specifically, in the bootstrapping process to calculate a simulated brightfield image to compare with the measured image, the radial distribution of the brightfield image is calculated by normal binning (histogram) compared to convolution with a Gaussian kernel. This choice was made to keep the calculation times manageable, and optimizing of this procedure, possibly through a two step (course and fine) optimization process is expected to yield improved results.

The STORM optimization results (lower row, Figure 7) do closely match the ground-truth result. Any deviation is mostly originating from a small number of cells which do not converge to the correct solution. Since the population is larger for lower photon brightfield images, and thus less accurate segmentation, this is be attributed to poor initial guesses for the coordinate system.

Finally, using an optimizer that is restricted by bounds on the optimization parameters is likely to reduce the subpopulations in both brightfield and STORM optimization. These features and the issues mentioned above will be addressed in a future performance update of ColiCoords.

## Conclusions

ColiCoords is an open-source software package to analyse fluorescence microscopy data of rod-shaped cells. It allows for the transformation of Cartesian coordinates from any data source to cellular coordinates. This transformation can then be used to obtain distributions of fluorescence along the cell long or short axes or its perimeter and align the whole cell to allow for the combination of data from many cells.

The open source nature and Jupyter-notebook based workflow together with an open and compact file format promotes open and reproducible analysis of microscopy data. As illustrated in this paper, ColiCoords facilitates the publication in the form a of a ‘reproducible article’(88), as recently featured eLife, where data and code to generate the figures are bundled together with the publication. The code and data to generate the figures in this article is available online (see Methods).

ColiCoords’ code is well documented and the main features are illustrated by example notebooks, which can be tested live on Binder. The software itself can be installed directly from both the Anaconda and PyPi package managers.

Analysis of synthetic data has shown that we can determine the cellular coordinates of SMLM super-resolution datapoints based on only the brightfield image of the cell with high accuracy (5%). If the coordinate system is derived from super-resolution measurements of a membrane marker, the accuracy is further increased (1%).

Although the current state of ColiCoords is a mature project, future updates are planned to be released, either by us or in collaboration with the community. Three major updates are planned; first a performance update to increase speed and accuracy of the optimization process, second a metadata update allowing the storage of image-associated metadata, and finally an update to allow the organization of cell objects into a hierarchical structure to process lineage and temporal information.

## Methods

### Cy3B-NHS staining

*E. coli* MG1655 cells were grown overnight in LB medium and subsequently diluted 1000 times in EZ rich medium (Teknova) with 0.2% glucose. The cells were grown until the culture reached an OD600 value of 0.3 after which the cells were centrifuged at 3000 rcf for 5 minutes and resuspended in PBS buffer. The amino-reactive dye Cy3B-NHS (GE Healthcare) was added to a final concentration of 1 mM and incubated while protected from light at room temperature for 1 hour(89). Next, the cells were centrifuged at 3000 rcf for 10 minutes and the cell pellet was resuspended in PBS.

### eGFP staining

*E. coli* BL21D3 cells were grown overnight in LB medium and subsequently diluted 1000 times 1000 times in EZ rich medium (Teknova) with 0.2% glucose. The growth media are supplemented with the appropriate antibiotics. When the cell culture reached an OD600 of 0.3 eGFP expression was induced with IPTG at a final concentration of 200 uM for 30 minutes at 37 °C.

For imaging, 3 *µ*L of the bacterial culture was transferred to a microscope coverslip and covered by an agarose pad(90, 91). Imaging was done on an Olympus IX83 inverted microscope with an Olympus UAPON 100× NA 1.49 TIRF oil immersion objective. The excitation light (514 nm, Coherent) was coupled into the objective via the ET - 442/514/561 Laser Triple band set (69904, Chroma) and the fluorescence was collected on a electron multiplying charge-coupled CCD camera (512×512 pixel, C9100-13, Hamamatsu). To focal position was held constant by the Olympus ZDC2 and images on multiple positions were collected automatically using Olympus’ CellSens software.

### Super-resolution imaging

LacY-eYFP-expressing *E. coli* C41 cells were grown overnight in LB medium with appropriate antibiotics and subsequently diluted 1000 times in EZ rich medium (Teknova) with 0.4% glycerol. When the cell culture reached an OD600 of 0.3, Lacy-eYFP expression was induced with 0.01%(v) L-arabinose for 30 minutes at 37 °C.

The cells were centrifuged at 3000 rcf for 5 minutes and resuspended in EZ rich medium with 0.2% glucose. 3 *µ*L of the bacterial culture was transferred onto a microscope coverslide and covered by an agarose gel pad.

Images were captured with an exposure time of 50 ms using the open source software *µ*Manager(92, 93).

To obtain the final STORM reconstruction 2000 frames were collected which were processed with the ImageJ plugin ThunderSTORM(60).

All data and code to used generate the figures in this article can be found at Zenodo (DOI: 10.5281/zenodo.2637790) and GitHub, respectively.

## ACKNOWLEDGEMENTS

This work was financed by an ERC Starting Grant (ERC-STG638536 – SM-IMPORT to T. C.) and an ERC Advanced Grant (ERC-ADG694610–SUPRABIOTICS to A.H). We thank T. Economou for critically reading the manuscript, M. Roelfs for insightful discussions on fitting best practises and R. Wind for logo design.

